# Spinal Column Architecture of the Flexible SPP1 Bacteriophage Tail Tube

**DOI:** 10.1101/2020.06.23.166439

**Authors:** Maximilian Zinke, Katrin A. A. Sachowsky, Carl Öster, Sophie Zinn-Justin, Raimond B.G. Ravelli, Gunnar F. Schröder, Michael Habeck, Adam Lange

## Abstract

Phage therapy has recently regained attention at combating multidrug-resistant bacteria. In 2019, tailed bacteriophages of the *Siphoviridae* family were engineered to successfully treat a disseminated bacterial infection after all other drugs had failed.(*1*) This family of phages features a long, flexible, non-contractile tail that has been difficult to characterize structurally. Here, we present the atomic structure of the tail-tube of the bacteriophage SPP1 – a member of this family. Our hybrid structure is based on the integration of structural restraints from solid-state NMR and a density map from cryo-EM. We show that the tail tube protein (TTP) gp17.1 organizes into hexameric rings that are stacked by flexible linker domains and, thus, form a hollow flexible tube with a negatively charged lumen suitable for the transport of DNA.

**One sentence summary:** Integrative structural biology by solid-state NMR and cryo-EM enables structure determination of the flexible tail of the bacteriophage SPP1.

## Main Text

Tailed bacteriophages – the order of *Caudovirales* – comprise the prevailing majority of known phages and are subdivided into three families based on their tail morphology. *Podoviridae* feature a short tail, *Myoviridae* a long, contractile tail and *Siphoviridae* a long non-contractile, flexible tail, respectively(*2*). The latter two possess a helical tail tube assembled around a tape measure protein that is tapered by a tail completion protein. Contractile tail tubes are furthermore environed by a sheath. These tail-structures are crucial for host-cell recognition, membrane penetration and DNA transport into the host. Recently, a high-resolution cryo-EM structure of the short fiber-less tail from the *Podoviridae* T7 phage was reported at a resolution of 3.3 Å.(*3*) Also, a cryo-EM structure of the pre-host attachment baseplate including two rings of the tail tube and sheath proteins from the *Myoviridae* T4 phage was solved at a resolution of 3.8-4.1 Å(*4*), and a cryo-EM reconstruction focused solely on the tail tube was obtained from the same images at a resolution of 3.4 Å(*5*). Structural analysis of the tube arrangement of *Siphoviridae* phages was for long hindered by the variable tail bending that results from its flexibility (see Figure 1 of Tavares et al.(*6*)). Structural information was limited to pseudo-atomic models, which were generated for phages λ(*7*) and SPP1(*8*) based on solution NMR structures of monomeric tail tube proteins (TTPs) and for the T5 phage by fitting X-ray structures of monomeric TTPs into a 6 Å cryo-EM density map(*9*). In 2020, a cryo-EM model of the baseplate of the *Staphylococcus aureus* 80α phage was reported, which includes two rings of the tail tube that are anchored within the baseplate and are, thus, not part of the flexible tube region.(*10*) All models show a striking structural homology between TTPs from *Myoviridae and Siphoviridae*; as well as between these phage TTPs and the tube-forming proteins from other injection systems, like the bacterial type VI secretion system(*11*) and the extracellular injection system from bacteria and archeae(*12*): The tube-forming proteins share a common fold composed of two orthogonally packed β-sheets that hexamerize through the formation of an inner β-barrel that defines the lumen of the tube. However, variable elements such as loops, N- and C-arms are critical to mediate intermonomer contacts driving tube assembly in these systems(*7*–*9*), and very sparse structural data are available on the position of these elements within the long-tailed phage tubes.

**Fig. 1.**
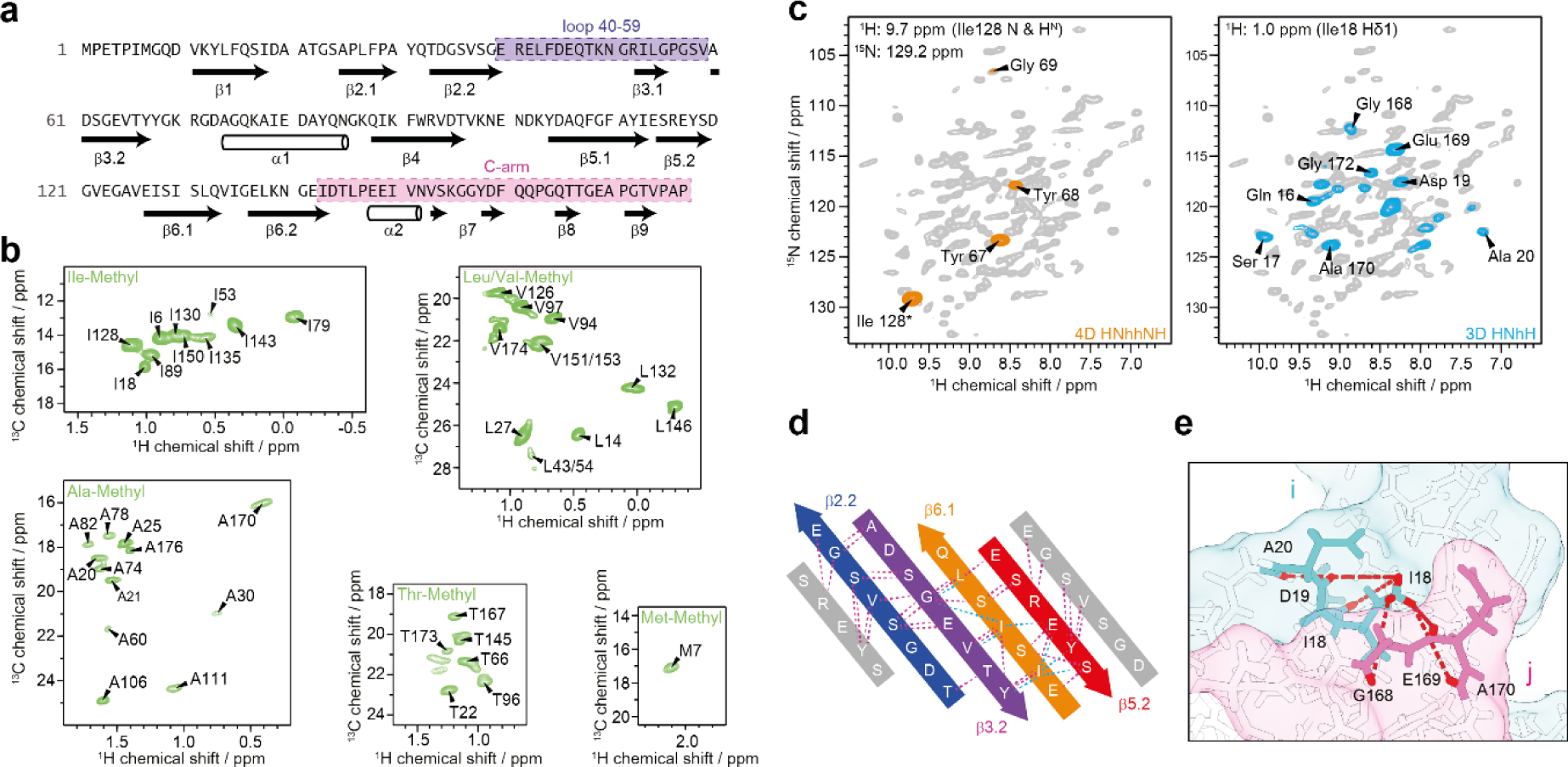
Solid-state NMR data for hybrid structure calculation defines local structure. **a**, Backbone dihedral angles define the secondary structure. **b**, Isoleucine Cδ1-, alanine Cβ-, leucine/valine Cδ/Cγ-, threonine Cγ2-, and methionine Cε-methyl labeling renders those moieties “NMR-visible” and yields highly resolved NMR spectra. **c**, Long-range restraints between amide protons are extracted from a 4D HNhhNH spectrum, whereas long-range restraints between amide and methyl groups are extracted from a series of 3D HNhH spectra. 2D planes from both types of spectra are superimposed on a 2D hNH fingerprint spectrum (grey). **d**, Representative long-range restraints defining the inner β-barrel of the tail tube. The colored β-strands belong to one monomer, whereas the grey β-strands are from the neighboring subunits. Amide-amide contacts are highlighted in violet, amide-methyl contacts in cyan. **e**, Schematic representation of long-range distance restraints between the Cδ1 methyl group of Ile18 and amide groups, defining the interface between the N-terminus of one monomer i (including Ile18; in cyan) and the C-arm (143-176) of another monomer j (in pink) within the tail tube. Protons are colored in red.

In the *Siphoviridae* SPP1 phage, the tail tube consists of the TTPs gp17.1 and gp17.1* in a ratio of 3:1, with the latter being generated by a translational frameshift adding a fibronectin type III (FN3) domain to the C-terminus of the protein. However, virions only containing gp17.1 are still viable and infectious, indicating that the additional C-terminal FN3 domain is dispensable for phage assembly and infection.(*13*) gp17.1 monomers are unstable in solution and spontaneously self-polymerize into long tubes *in vitro* which are indistinguishable from native tubes.(*8, 14*) Previously, we presented the proton-detected solid-state NMR (ssNMR) assignment of deuterated, 100% back-exchanged gp17.1 tubes and deduced secondary structure information from the assigned chemical shifts, which confirmed and extended an existing homology model of a polymerized gp17.1 subunit.(*8, 14*) Additionally, we introduced new concepts based on the use of specifically labeled isoleucine-methyl groups to simplify ssNMR data, inspired by earlier progress based on methyl-labeling in solution state NMR.(*15, 16*). This allowed for the collection of unambiguous long-range distance restraints within and between subunits of the tail tube.(*17*)

However, due to the large size and inherent heterogeneity of this system the amount of data collected from these experiments does not suffice for a confident structure calculation. Therefore, we set out to expand the labeling strategy for long-range distance restraints to further methyl groups, as well as to integrate the NMR data with a 3.5 - 6 Å cryo-EM map for hybrid structure calculation. Solid-state NMR is a powerful method to study the structure and dynamics(*18*) of insoluble proteins, such as amyloid fibrils(*19, 20*) or supramolecular assemblies(*21*–*24*). An integrated structure calculation approach(*25*) in combination with cryo-EM has proven highly successful in previous applications by us(*26*) and others(*27*–*32*). The complementarity of solid-state NMR and cryo-EM can also be appreciated by work on bactofilin cytoskeletal filaments.(*33, 34*)

### Hybrid structure calculation

To determine the structure of the tail-tube of SPP1, we performed a hybrid structure calculation using the Inferential structure determination (ISD) approach(*35*) integrating data from solid-state NMR and cryo-EM simultaneously. During structure calculation only the structure of a single monomer was represented and refined. The structures of the other subunits were generated by applying symmetry operators to the subunit structure. The use of an exact symmetry is justified by the NMR data that are only consistent with a highly symmetric sample. We represented two stacked rings in the structure calculation. Each ring was composed of six subunits such that interactions between twelve subunits were considered in the structure calculation.

For the collection of solid-state NMR long-range distance restraints, we overexpressed a set of differently labeled TTP gp17.1 in *E. coli*, purified them, and let them self-polymerize into native-like tail tubes as detailed in the Methods. Torsion angles were predicted based on assigned backbone chemical shifts (Figure 1a). Specific precursor molecules were supplemented during protein expression in deuterated media, introducing NMR visible methyl groups within certain amino acids of gp17.1 (Figure 1b). We produced samples that were homogeneously methyl and ^15^N labeled, as well as samples that were heterogeneous mixtures of 50% methyl-labeled and 50% ^15^N labeled subunits. The latter samples were used to detect intermolecular interfaces, as previously described by us.(*17*) All eleven investigated methyl-labeled and/or deuterated samples and their precursors are listed in Table S1. 4D and 3D proton-detected ssNMR experiments at 40 kHz magic-angle spinning (MAS) and 900 MHz proton Larmor frequency were used to probe long-range distance restraints between the following: 1.) amide groups (Figure 1c, left panel)(*36, 37*); 2.) methyl groups; 3.) methyl and amide groups globally (Figure 1c, right panel); 4.) methyl and amide groups at protein-protein interfaces. All of these experiments generated highly unambiguous restraints due to their high-dimensionality (4D) or spectral simplicity (amino-acid specific methyl labeling) – as visualized in Figure 1d where a set of consistent restraints (amide-amide and methyl-amide contacts) defines the inner β-barrel motif of the tail tube formed by the β-strands β2.2, β3.2, β6.1 and β5.2. In mixed labeled samples (as detailed in Table S1) magnetization transfer between methyl and amide groups is solely possible at protein-protein interfaces. Hence, these samples deliver a set of restraints that defines the organization of gp17.1 subunits within the tube. Figure 1e shows an exemplary protein interface between the N-terminus of a subunit, including the Ile18 labeled methyl group, and the C-terminus of another subunit.

For cryo-EM experiments, we purified ΔN-3 gp17.1 which is indistinguishable from wt gp17.1 as judged by solid-state NMR (See Figure S1). Curvy tubes were observed in the micrographs (Figure S2). For image processing those tubes that appeared most straight were selected. 3D reconstruction (see Methods) yielded a density map with an average resolution of 4 Å (Figure 2a). The local resolution varies significantly (Figure 2b-d), from about 3.5 Å at the inner ring where the β-strands are well resolved (Figure 2e-f), to worse than 5 Å at the periphery.

**Fig. 2.**
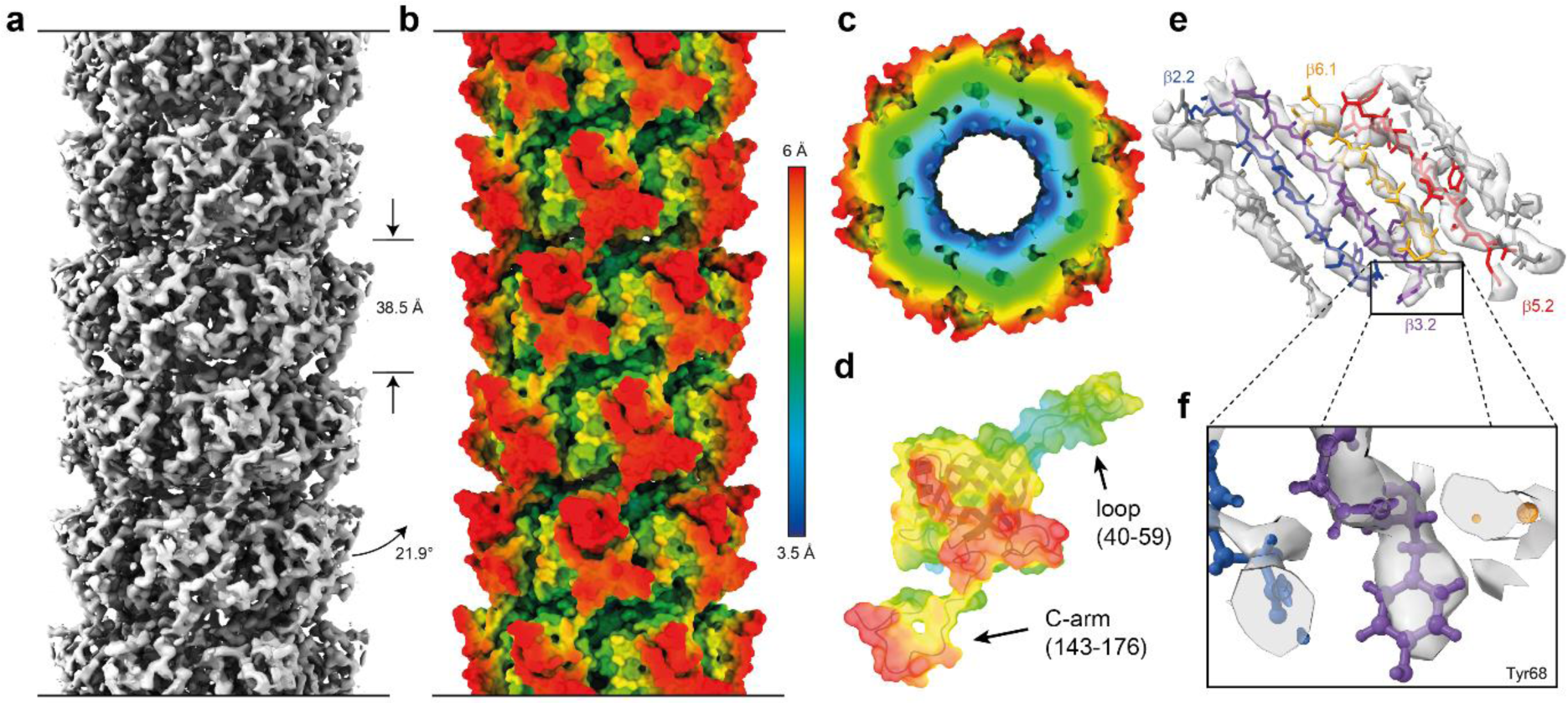
Cryo-EM data for hybrid structure calculation provides global and local information. **a**, 3.5 - 6 Å cryo-EM map of the tail tube of SPP1 consisting of polymerized gp17.1 subunits does not only allow to deduce symmetry restraints but also limits the position of all atoms within the density. **b-d**, Local resolution of the cryo-EM map increases going from outer to the inner surface of the tail tube. **e**, The inner region of the map reveals a highly resolved β-barrel and allows for the positioning of bulky sidechains as exemplified for Tyr68 (**f**).

### Overall structure of the SPP1 tail tube

Figure 3 shows the structure of the tail-tube of gp17.1 as determined by hybrid structure calculation (see also Movie S1 and Figure S3) with a heavy atom RMSD of 1.8 ± 0.9 Å (Backbone RMSD of 1.1 ± 0.5 Å) over the entire protein sequence. The monomer of gp17.1 forms a central β-sandwich-type fold consisting of eight β-strands. This fold is flanked by an α-helix (74-86) and loop regions, of which the long C-terminal arm (C-arm, 143-176) stands out (Figure 3a). Six gp17.1 subunits assemble into a ring with the inner 24 β-strands forming a β-barrel that defines the inner lumen of the tube. This inner lumen exhibits a negative electrostatic potential (see Figure S4) which facilitates sliding of the viral DNA through the tube by repelling it from the surface. Additionally, inner ring contacts are mediated by the large loop region 40-59. The α-helices are arranged almost parallel to the tail tube axis (Figure 3b). These rings stack onto each other with a rotation of 21.9° forming a right-handed helical, hollow tube (Figure 3c). The surface of one gp17.1 subunit features a hydrophobic patch that is shaped by sidechains of various hydrophobic amino acids (Figure 3d). In the context of the tail tube complex the C-arm of the superjacent subunit folds onto the outer β-sheet of the β-sandwich fold by anchoring the sidechain of Gln162 into a pocket (Figure 3e). This interaction obscures the lipophilic area whereupon the complex is stabilized, because the number of unfavorable hydrophobic contacts with the solvent is reduced (Figure 3f). This explains why a previously reported C-terminally truncated mutant of gp17.1 remains monomeric(*17*). Additional ring-to-ring contacts are mediated by the loop region 40-59 which interacts with five neighboring subunits - mostly by establishing electrostatic contacts (Figure 3g).

**Fig. 3.**
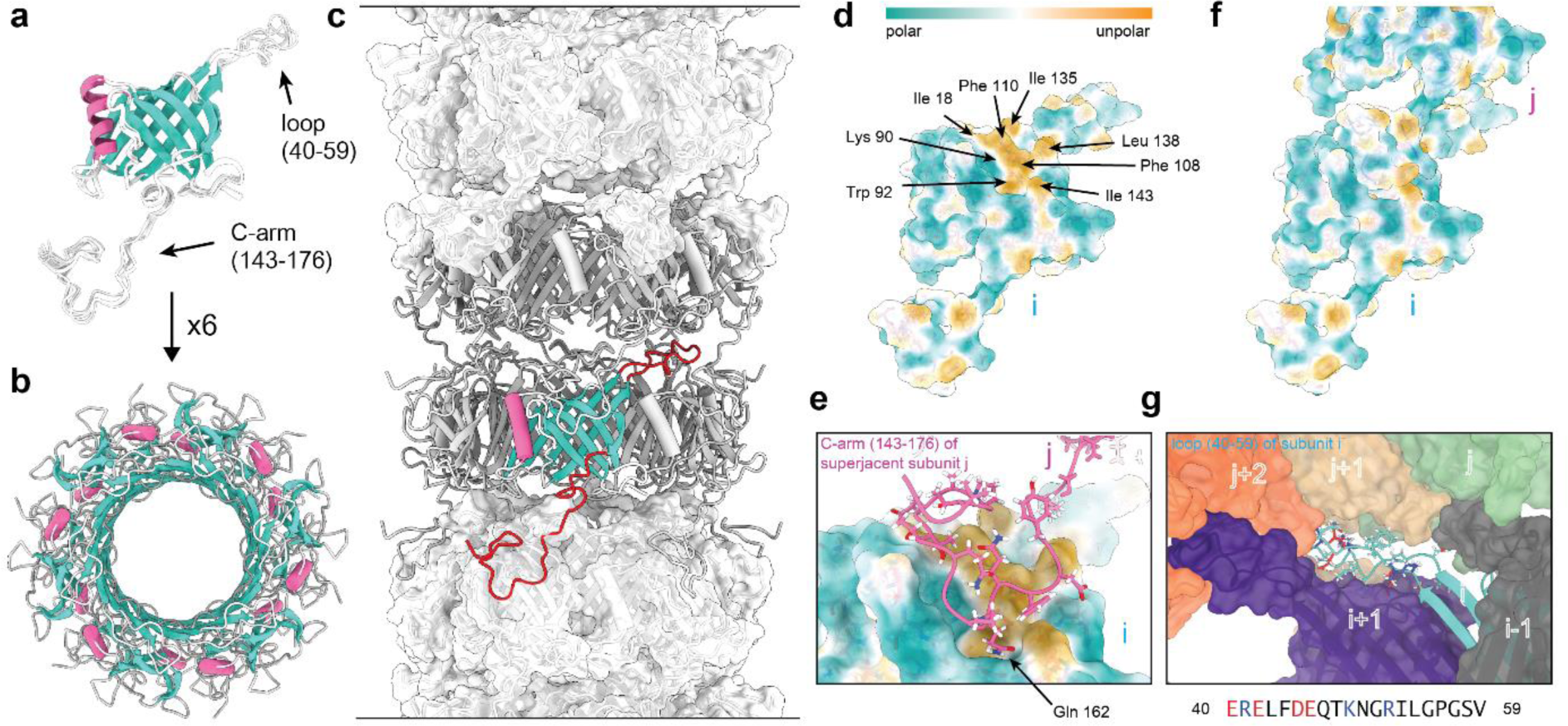
Structure of polymerized gp17.1 forming the tail tube of the bacteriophage SPP1. **a**, Final ten lowest-energy structures of a gp17.1 subunit which consist of a central β-sandwich-type fold that is flanked by an α-helix, a large loop and an extended C-terminal arm (C-arm). **b**, Six gp17.1 monomers form a hexameric ring. The inner β-sheets of the β-sandwiches organize in a β-barrel motif that forms the lumen of the tube. **c**, These hexameric rings stack onto each other in a helical fashion creating a hollow tube. Ring-to-ring contacts are mediated by the two loop regions (highlighted in red) – especially by the C-arm that folds onto the subjacent ring. **d**, The molecular lipophilicity potential of gp17.1 reveals a hydrophobic patch on the surface of one subunit i. **e-f**, This unpolar area is obscured by the C-arm of the superjacent subunit j within the complex of the tail-tube – by anchoring the sidechain of Gln162 into a pocket. **g**, The loop of subunit i features mostly electrostatic interactions with five neighboring subunits within the complex. Charged amino acids are colored in red (negative) and blue (positive).

Structural alignments of gp17.1 and existing TTP structures of other systems show high similarity as expected (Figure S5) – all featuring hexameric, helically stacked rings with subunits consisting of a β-sandwich-type fold and one parallel α-helix. Loop 40-59 is also present at the interface between subunits in the phages 80α(*10*) and T4(*5*), suggesting that it is a conserved structural element across both *Siphoviridae* and *Myoviridae* families. The mentioned C-arm is not observed in the *Myoviridae* T4, however it is present in 80α(*10*) (even if not completely resolved). It might be a critical element regulating the tail structure in a subgroup of *Siphoviridae*. The *Myoviridae* T4(*5*) phage TTP gp19 features two additional linkers that mediate intermolecular interactions – which facilitates contact to 10 different subunits over an area of 6706 Å^2^ within the tail tube. gp17.1, however, only interconnects with 6 different subunits over 4850 Å^2^. This dramatically reduced contact – in addition to not being bundled in a sheath - is expected to enable flexibility of this *Siphoviridae* tail tube.

### Dynamic regions mediate tail bending

To determine the driving forces contributing to the flexibility of the tail tube of SPP1, we created a model of a bent tail tube based on the structure of monomeric gp17.1 and 2D class averages of bent tubes as detailed in the Methods. As shown in Figure 4a (and Movie S2), most structural changes during the bending process happen on the outer edge of the curve. This suggests that the bending of the tube is facilitated by stretching of certain linker regions (as opposed to compression). The hinge regions required for the structural reorganization from a straight to a bent state (Figure 4b) match regions in the cryo-EM density with pronounced variances (Figure 4c). These areas comprise regions forming inter-ring contacts – especially the C-arm (143-176), its binding interface on the subjacent subunit and the loop (40-59).

**Fig. 4.**
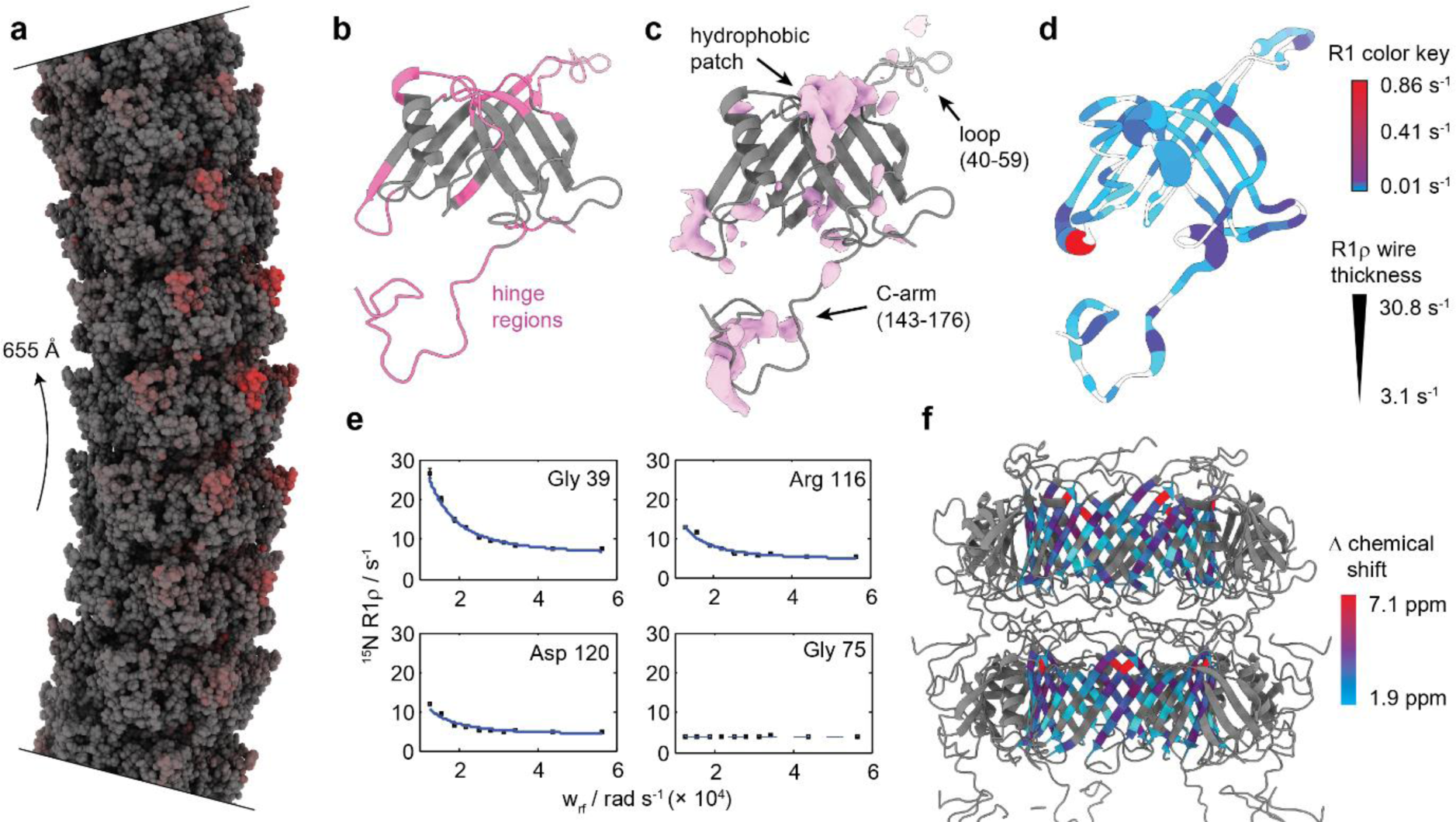
Bending of the tail tube is mediated by flexible hinge regions. **a**, Model of a bent SPP1 tail tube with a curvature radius of 655 Å. The model is based on the structure of straight tubes and 2D class averages of bent tubes. Most structural changes are found on the outside of the curve (red) which implies that the bending process is mediated by stretching. **b**, Regions that act as hinges during bending of the tube are colored in pink. **c**, Variances in the cryo-EM map (pink) match the hinge regions. **d**, Also, hinge regions are associated with highest ^15^N R_1_ (color key) and R_1ρ_ (thickness of wire) relaxation rates. White coloring represents missing values. **e**, Decaying relaxation dispersion profiles indicate the presence of slow motions. **f**, Relaxation dispersion profiles of the inner β-barrel can be fitted in a correlated manner to a two-state exchange process. Smallest chemical shift changes correlate with middle regions of the β-barrel which are furthest away from the hinge regions (color key).

Additionally, we analyzed ^15^N R_1_ and ^15^N R_1ρ_ relaxation rates of fully polymerized gp17.1 by solid-state NMR as detailed in the Methods (Figure S6). R_1_ relaxation rates are sensitive to motions on the nanosecond timescale, whereas R_1ρ_ rates report on motions on the nanosecond to millisecond timescale.(*38*) Figure 4d shows both values mapped onto a gp17.1 subunit within the tail complex. The inner β-barrel shows nearly no motion on these timescales - which correlates with these areas being highest resolved in the cryo-EM density. On the opposite, the hinge regions are associated with the highest relaxation rates, demonstrating that these areas are highly dynamic. Furthermore, we investigated ^15^N relaxation dispersion which is sensitive to motions on the millisecond timescale (Figure S7-S9). Figure 4e shows representative ^15^N relaxation dispersion curves; flat profiles indicate the absence of motion, whereas decaying profiles indicate the presence of dynamics. Most residues are involved in slow motions (Figure S10). However, only residues belonging to the β-barrel can be fitted in a combined approach to a two-state model indicating the existence of two distinct conformations of the β-barrel (Figure 4f). Calculated chemical shift differences between both states are higher for residues in proximity to the hinge regions. Thus, we propose that these two states could represent the two extrema of tube bending – either lying on the inside or the outside of the curve. The global, collective nature of this motion does not impose heterogeneity onto the cryo-EM map since only straight tubes are considered for structure calculation.

Overall, our hybrid data support a model where the C-arm (143-176) and the loop (40-59) act as bellows contributing to tail tube bending by stretching. Our dynamic structure of the tail tube is reminiscent of a molecular spinal column. The hexameric rings forming the inner β-barrel would be in this picture the vertebrae, while the flexible parts (C-arm and loop) correspond to the intervertebral discs. The flexibility of the system might facilitate the screening of the bacterial membrane to find the receptor for infection initiation. We expect that the combination of sophisticated ssNMR experiments and cryo-EM will help to characterize structures of other dynamic and/or flexible supramolecular assemblies, in particular those systems where the conformational flexibility leads to a lack of resolution in cryo-EM reconstructions – while depending on the timescale of these dynamics the quality of NMR spectra may not be affected.

## Acknowledgements

We thank Dr. Paulo Tavares for valuable discussions and Dr. Sascha Lange for help with the sample production. Molecular graphics and analyses were performed with UCSF ChimeraX, developed by the Resource for Biocomputing, Visualization, and Informatics at the University of California, San Francisco, with support from National Institutes of Health R01-GM129325 and the Office of Cyber Infrastructure and Computational Biology, National Institute of Allergy and Infectious Diseases.

## Author contributions

M.Z. produced samples for NMR and cryo-EM measurements and performed solid-state NMR measurements. M.Z., S.Z.-J., and A.L. analyzed data from solid-state NMR measurements. C.O. and M.Z. analyzed the relaxation data. K.A.A.S., R.R., and G.F.S. performed cryo-EM measurements, K.A.A.S. and G.F.S. analyzed cryo-EM data, M.H. performed the hybrid structure calculation, A.L., M.H., and G.F.S. conceived this study, M.Z., G.F.S., M.H., and A.L. wrote the manuscript.

## Data and materials availability

Solid-state NMR chemical shift assignment is deposited at the Biological Magnetic Resonance Data Bank under the accession code 27468. Cryo EM electron density map is deposited at the Electron Microscopy Data Bank under the accession code EMD-10792. Protein structures are deposited at the Protein Data Bank under the accession codes 6YEG (Hybrid structure of the SPP1 tail tube by solid-state NMR and cryo EM - Final EM Refinement) and 6YQ5 (Hybrid structure of the SPP1 tail tube by solid-state NMR and cryo EM – NMR ensemble).

## Supplementary Materials

Materials and Methods

Figures S1-S13

Tables S1-S11

Movies S1-S2

References (39-61)

