## Supplementary material for "Spinal Column Architecture of the Flexible SPP1 Bacteriophage Tail Tube": SI

**This PDF file includes:**

Materials and Methods

Figs. S1 to S13

Tables S1 to S11

Captions for Movies S1 to S2

**Other Supplementary Materials for this manuscript include the following:**

Movies S1 to S2

### Materials and Methods

#### Protein sequences of gp17.1 used in this study

```
>gp17.1:

MGMPETPIMGQDVKYLFQSIDAATGSAPLFPAYQTDGSVSGERELFDEQTKNGRILGPGSVADSGEVTYYGKRGDAG

QKAIEDAYQNGKQIKFWRVDTVKNENDKYDAQFGFAYIESREYSDGVEGAVEISISLQVIGELKNGEIDTLPEEIVN

VSKGGYDFQQPGQTTGEAPGTVPAPHHHHHH
```

```
>ΔN-3 gp17.1:

MPIMGQDVKYLFQSIDAATGSAPLFPAYQTDGSVSGERELFDEQTKNGRILGPGSVADSGEVTYYGKRGDAGQKAIE

DAYQNGKQIKFWRVDTVKNENDKYDAQFGFAYIESREYSDGVEGAVEISISLQVIGELKNGEIDTLPEEIVNVSKGG

YDFQQPGQTTGEAPGTVPAPHHHHHH
```

#### Preparation of deuterated protein samples for proton-detected solid-state NMR measurements

Protein samples and their preparation are summarized in Table S1. In general, gp17.1 protein was expressed, purified and polymerized as described earlier(*14*). For methyl-labeling, $^{12}C_6$,$D_7$-glucose was used instead of $^{13}C_6$,$D_7$-glucose. For mixed-labeling, two differently labeled samples, e.g. 50% methyl-labeled and 50% $^{15}N$-labeled, were produced and mixed before polymerization following the procedure described by us before(*17*). Labile protons were 100% back-exchanged in all samples. The protein pellets, a few DSS crystals for spectral referencing and temperature control, and 1 µL of $D_2O$ for field locking were filled into 1.9 mm rotors provided with bottom spacers.

#### Preparation of fully protonated protein samples for carbon-detected solid-state NMR and cryo-EM measurements

gp17.1 and ΔN-3 gp17.1 were expressed, purified and polymerized as described earlier(*14*) ($D_2O$ was exchanged to $H_2O$). The protein pellets and a few DSS crystals for spectral referencing and temperature control were filled into 3.2 mm rotors. For cryogenic electron microscopy (cryo-EM) 20 mM sodium phosphate in the final buffer was replaced by 20 mM Tris-HCl (pH 7.4).

#### Solid-state NMR spectroscopy

Solid-state NMR spectroscopy of the methyl-labeled and/or deuterated protein samples was conducted with a 1.9 mm, four-channel ($^{1}H$, $^{13}C$, $^{15}N$, $^{2}H$) probe at 40 kHz magic-angle spinning (MAS) frequency and an external magnetic field strength according to 900 MHz $^{1}H$ Larmor frequency. The temperature was calibrated to around +18°C by means of internally added DSS. 2D hCH and 3D HNhH spectra were recorded as described previously(*17*), a 2D hNH spectrum was recorded as detailed before(*14*). Pulse program, acquisition, processing and reconstruction parameters for the 2D hCH, 3D HNhH, 3D HChH, and 4D HNhhNH spectra are summarized in Supporting Information Tables S2-S6.

Solid-state NMR spectroscopy of the fully-protonated samples was conducted with a 3.2 mm, triple-channel ($^{1}H$, $^{13}C$, $^{15}N$) probe at 11 kHz MAS frequency and an external magnetic field strength according to 900 MHz $^{1}H$ Larmor frequency. The temperature was calibrated to around +10°C by means of internally added DSS. 2D $^{13}C$-$^{13}C$ correlation spectra with 50 ms proton-driven spin diffusion (PDSD) mixing were recorded as fingerprints to compare wild-type gp17.1 with the ΔN-3 gp17.1 mutant.

Long-range distance restraints were extracted from the recorded spectra by peak picking in CcpNmr(*39*). Methyl groups were assigned based on our previous work(*17*), on the basis of the assignment precursor (Table S1) or by correlations to sequential amide groups. In the alanine-methyl sample (Table S1) protons scrambled into the Hγ2 position of isoleucines.

#### Relaxation measurements by solid-state NMR

$^{15}N$ $R_1$ and $^{15}N$ $R_{1\rho}$ relaxation rates were measured by a series of pseudo-3D experiments ($^{1}H$, $^{15}N$, delay/spinlock strength). The delay times for the $R_1$ pseudo-3D experiments were 0.5 s, 1 s, 1.5 s, 2 s, 3 s, 4 s, 8 s, 16 s and 32 s. For the $R_{1\rho}$ pseudo-3D experiments spinlock strengths of 9 kHz, 7 kHz, 5.5 kHz, 5 kHz, 4.5 kHz, 4 kHz, 3.5 kHz, 3 kHz, 2.5 kHz and 2 kHz and spinlock durations of 1 ms, 5 ms, 10 ms, 20 ms, 40 ms, 80 ms, 100 ms, 140 ms and 200 ms were used. The peak heights from the resulting 2D hNH correlation spectra were extracted with CcpNmr(*39*) and fitted as a function of relaxation time to a monoexponential function. This results in one global $R_1$ relaxation rate for each - in the 2D hNH spectra distinguishable - amide nitrogen and various spinlock strength-dependent $R_{1\rho}$ relaxation rates. For relaxation dispersion analysis the $R_{1\rho}$ values are plotted against the spinlock strength and fitted to a two-site Bloch-McConell equation(*40*):

$$R_{1\rho} = R_{1\rho,0} + \frac{p_{\mathrm{A}} p_{\mathrm{B}} \Delta\delta^2 k_{ex}}{{\omega_1}^2 + {k_{ex}}^2} = R_{1\rho,0} + \frac{\varphi_{ex} k_{ex}}{{\omega_1}^2 + {k_{ex}}^2} \quad (1)$$

For all resolved residues that are part of the β-barrel forming the inner lumen of the tail tube a combined fit was conducted with a single $k_{ex}$ coefficient for all residues and individual $\varphi_{ex}$ and $R_{1\rho,0}$ for each residue (See Table S7). The best fit was achieved by minimization of the target function $\chi^2$ as demonstrated previously(*41*). On-resonance

$R_{1\rho}$ relaxation rates were deviated from the observed $R_{1\rho,obs}$ relaxation rates, $R_1$ relaxation rates and the angle between the spinlock offset frequency ($\omega_1$) and the chemical shift offset from that (Ω) in a residue specific manner:

$$R_{1\rho} = \frac{R_{1\rho,obs} - R_1 \cos^2 \theta}{\sin^2 \theta} \quad (2)$$

$$\theta = \tan^{-1} \frac{\omega_1}{\Omega} \quad (3)$$

The errors were estimated by Monte Carlo simulations. The fits were repeated 250 times with the $R_{1\rho}$ errors being multiplied by a random number between 0 and 1. The $R_{1\rho}$ errors were calculated in a similar way by performing the fits 1000 times using average noise from the spectra as input.

**Cryo-EM image acquisition**

Cryo-preparation was performed on glow-discharged holey carbon films (Quantifoil R 1.2/1.3, 300 mesh) using a Vitrobot (FEI). With 110,000-fold nominal magnification 855 micrographs have been recorded on a Tecnai Arctica electron microscope operating at 200 kV with a field emission gun using a Falcon III (FEI) direct electron detector in integrating mode directed by EPU data collection software (version 1.5). Each movie was composed of 20 fractions. Each fraction contained 6 frames, i.e. a total of 120 frames were recorded per micrograph. The sample was exposed for 3 s to a total dose of 70 $e^-$/$Å^2$. Applied underfocus values ranged between 0.2 and 1.7 µm. The pixel size was calibrated to 0.935 Å as calibrated using gold diffraction rings within the powerspectra of a cross grating grid (EMS, Hatfield). Details of data acquisition are summarized in Table S8.

**Cryo-EM image processing and helical reconstruction**

MotionCor2(*42*) was used for movie correction and CTF parameters were fitted with Gctf(*43*). All image processing was done using RELION 2.1(*44*). Fibrils were manually picked, and segments were extracted with an interbox distance of 10 % of the box sizes, which yielded 69,282 segments. Box sizes were chosen as 200 pixels. Segments from micrographs for which Gctf estimated a resolution worse than 5 Å were discarded, which left 64,760 segments. 2D classification was used to select those classes that show sufficient detail in the class averages (Figure S11). The initial model for 3D reconstruction was built with the Relion *relion_helical_toolbox* program with the --*simulate helix* option, which places spheres along a helix, using a rise of 40 Å and a twist of 21º.

The initial model was refined using 3D classification (K=3). The class yielding the highest resolution was used as initial model for further 3D classification with all particles using K=5 classes and a T value of 4. The best resolved class contained 10682 segments which were used for further refinement. A soft mask enclosing three rings was used for further refinements. The number of filaments and segments used for the final reconstruction was 1,866 and 5,965, respectively. The final optimized helical symmetry was C6, with a helical rise of 38.46 Å and a twist of 21.89º. Gold-standard refinements were performed by selecting entire fibrils and splitting the data set accordingly into an even and odd set. The Fourier shell correlation was computed between two half maps. To obtain a robust resolution estimate, the FSC curve was fitted using $1/[e^{((x-A)/B)}+1]^C$, yielding A=0.122, B=0.015, and C=0.228. According to the 0.143 criterion the obtained resolution is 4.0 Å (Figure S12). Image processing and reconstruction details can be found in Table S8. The final map was sharpened with the EMAN2 tool *e2proc3d.py* with a B-factor of -150 Å$^2$, locally normalized and filtered to 3.5 Å.

#### Structure calculation

The hybrid structure calculation aims to combine the experimental data from NMR and cryo-EM. Distance restraints were derived from NMR peak lists and incorporated using a logistic restraint potential similar as described before(*45*). The density map from cryo-EM was incorporated as a real-space map restraint.(*46*) Both data sets pose their own challenges: The NMR distance restraints are highly ambiguous due to the helical symmetry of the tail tube. NMR peaks stemming from the homogeneously mixed samples can result from a contact within the monomer or from contacts between different subunits (i.e. between the monomer and one of its virtual copies generated by the symmetry operators). However, NMR peaks stemming from the heterogeneously mixed samples can clearly be assigned as intermolecular long-range restraints. We considered all possible interactions between all members of both hexameric rings. Therefore, in total twelve possible contacts were combined as an ambiguous distance restraint. Another challenge is posed by the generous upper bound of 7 Å for the distance restraints. Also, the cryo-EM map itself has a substantial resolution inhomogeneity and is not sufficient for an unambiguous tracing of the backbone in the outer β-strands and the C-terminus.

A particular challenge was posed by the estimation of the registers of adjacent β-strands. Due to the insufficient resolution of the density map and the large distance upper bounds, many relative registers between adjacent strands seem possible in principle. To infer the register that is most consistent with the NMR and cryo-EM data, we developed a new probabilistic restraint that probes all possible registers between adjacent strands. The estimated registers were then imposed as additional hydrogen bonding restraints to increase the regularity of the gp17.1 tail-tube structure.

Only the combination of the local information provided by the NMR restraints with the global shape information encoded in the cryo-EM map allowed us to compute a near-atomic structure of the tail tube of gp17.1. For example, in the initial phase of the project we tried to compute a structure based only on the NMR restraints and homology information with literature values for the symmetry parameters. Although the β-sandwich and the α-helix can be computed from this information, it was not possible to model the loop (40-59) and the C-terminus correctly. Due to the ambiguity of the NMR restraints resulting from the helical symmetry it is not clear from the NMR data which subunit of the subjacent ring is contacted by the C-arm.

In the final stage of the hybrid structure calculation, we used an iterative approach in which hybrid structure calculation with ISD was alternated with a pure EM refinement using Coot(*47*) and Phenix(*48*). MDFF simulations for 5 ns were used on intermediate models to guide model building in Coot.

Finally, two different models were generated for final interpretation (and were also deposited to the PDB): One (ensemble of) models represents the ensemble from the final ISD refinement (PDB ID: 6YQ5). The other model (PDB ID: 6YEG) represents a standard refinement against the EM density (including cycles of Coot and Phenix) starting from the hybrid NMR-EM model obtained from ISD. The statistics of the solid-state NMR long-range distance restraints, the violations and the RMSDs of the final ensemble are summarized in Tables S9-11.

#### Variance map

To determine the structural variance in the dataset a bootstrapping analysis was performed. For this, 300 density maps were reconstructed (with fixed orientations and shifts) from randomly resampled (with replacement) sets of segment images, using the *relion_reconstruct* command with C6 and helical symmetry.

The density variance calculated directly over such resampled density maps often leads to artifacts (strong noisy variance outside the particle and strong variance at symmetry axes). We therefore computed instead the isosurface variance map, which yields a clearer view of the structural variance. To compute the isosurface variance, all 300 density maps were first low-pass filtered, and then masks were computed using a density threshold of 0.162. Then a Gaussian filter was applied to all 300 masks. Finally, the variance of all masks was computed, which yields the isosurface variance map (Figure 4 c). Since symmetry was used during the reconstruction, symmetry-breaking variance is not visible.

#### Model of a bent SPP1 tail tube

For the analysis of the curved filament regions, curved filaments were picked in short segments. The average number of segments per picked filament was only 4, whereas it was 13 for the straight filaments. 12,259 segments was obtained from the curved filaments, which were extract with a larger box size of 400 pixels to clearly see the curvature. From a 2D classification with 50 classes the best defined classes were chosen and yielded 2,735 segments. Further 3D classification yielded a final reconstruction with 1,418 particles at a resolution of only about 17 Å. Due to the curvature, no helical symmetry could be used.

On the basis of the 2D class averages of bent tail tubes (Figure S13) a curvature radius of 655 Å (inner radius = 655 - 63.3/2 = 623.4 Å; outer radius = 655 +63.3/2 = 686.7 Å), an inner distance between subunits of 38.5 Å, an outer distance between subunits of 47.0 Å and an angle between subunits of 3.5° could be determined. The distance between neighboring rings on the inside is the same as in the straight filament, however, the distance on the outside is larger. The curvature is therefore induced by stretching the outside while the inside distances remain unchanged compared to the straight filament. The curvature from the 2D class averages represents the average, most populated curvature. A maximum curvature of 560 Å could be extracted from the micrographs which results in a maximum angle between subunits of 4.2° (Figure S2). This geometric information was used to build a 10-ring bent tail tube from gp17.1 monomers. For this, copies of one ring from the straight tail tube were translated and rotated accordingly.

The rationale for building this bent model was to impose the observed curvature and ring distances, but to keep the local structure and subunit contacts as similar as possible to the straight tail tube, as we do not have high-resolution information on the curved tail tube. Therefore, a network of harmonic distance restraints (random atom pairs between 3 and 15 Å) was defined with target distances from the straight tail tube. In addition the β-sheets at the inside of the tubes were position-restrained to keep curvature and relative ring positions. DireX(*49*) was used to optimize the model under these distance and position restraints (without density map restraints). The curved model (Figure 4a) represents a model that is closest (in local structure and subunit contacts) to the straight tail tube, while adopting the imposed curvature and ring distances. The amount of fulfilled restraints after bending reveals regions of the protein that are exposed to environment changes (red color coding in Figure 4a).

ChimeraX(*50*) morph command was used to create a trajectory between the straight and the bent tail tube (standard settings). During that procedure hinge and core regions are identified by a reimplementation of the morph server(*51*).

**Figure creation**

Figures were created with ChimeraX.(*50*)

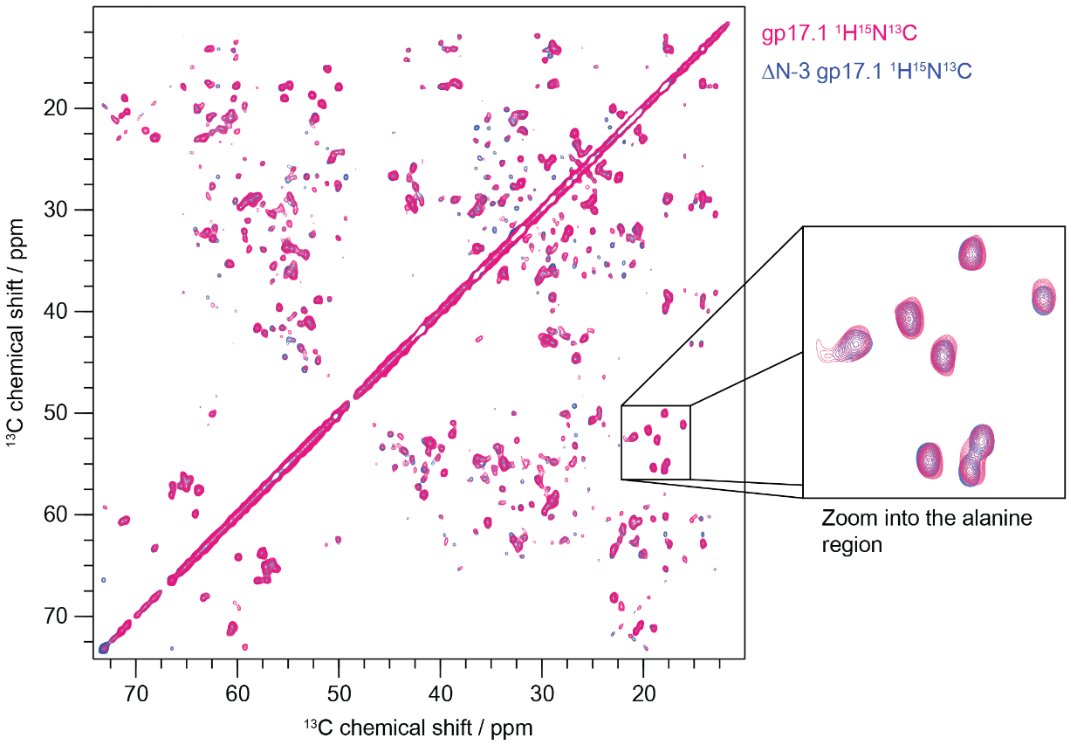

**Fig. S1.** Aliphatic region of 2D hCC solid-state NMR correlation spectra of fully-protonated gp17.1 and ΔN-3 gp17.1 tail tubes at 11 kHz MAS and 900 MHz external magnetic field strength. The zoom into the alanine region reveals that both spectra show the same fingerprint, demonstrating that the N-terminal truncation does not impair the structural organization of the tail tube.

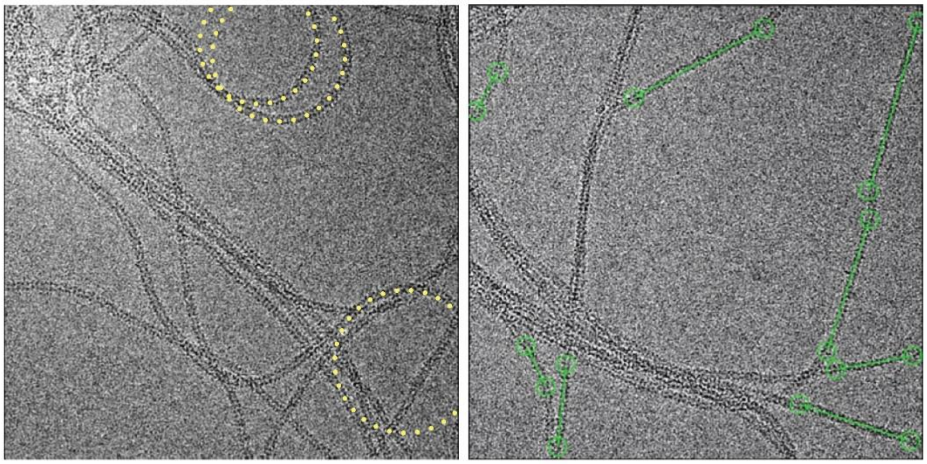

**Fig. S2.** Typical micrographs of polymerized gp17.1 tail tubes. a) The flexible tubes show variable bending and cross each other. The curvature radius of the tubes can be measured as indicated by the yellow circles. b) Exemplary picked straight filament segments for cryo-EM image processing. The green circles label the starting and ending coordinates.

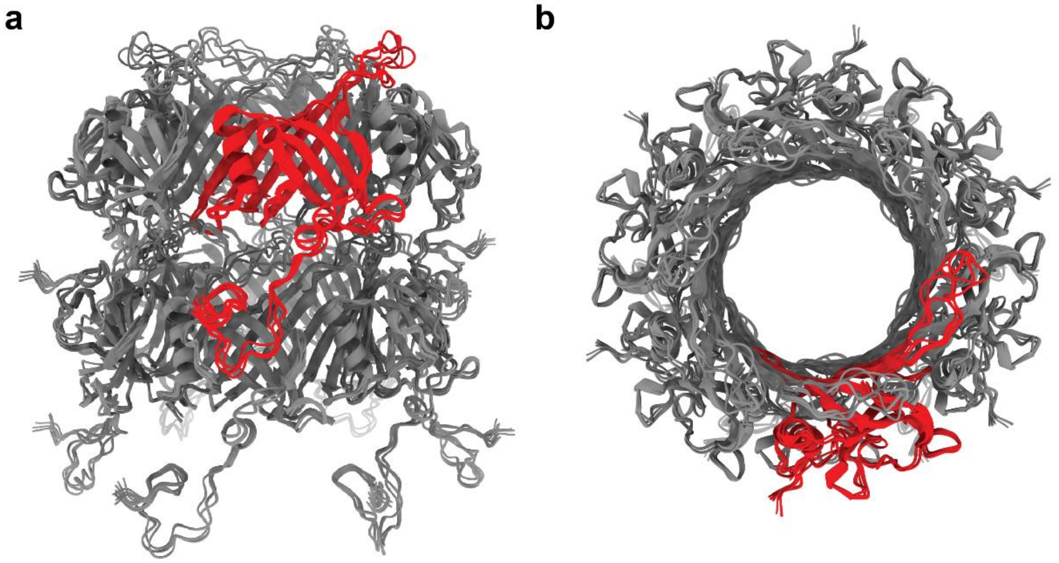

**Fig. S3.** Final ten lowest-energy structures of two SPP1 tail-tube rings after hybrid structure calculation (PDB ID 6YQ5). One gp17.1 subunit within the top ring is highlighted in red. The assembly is shown from the side a) and from the top b).

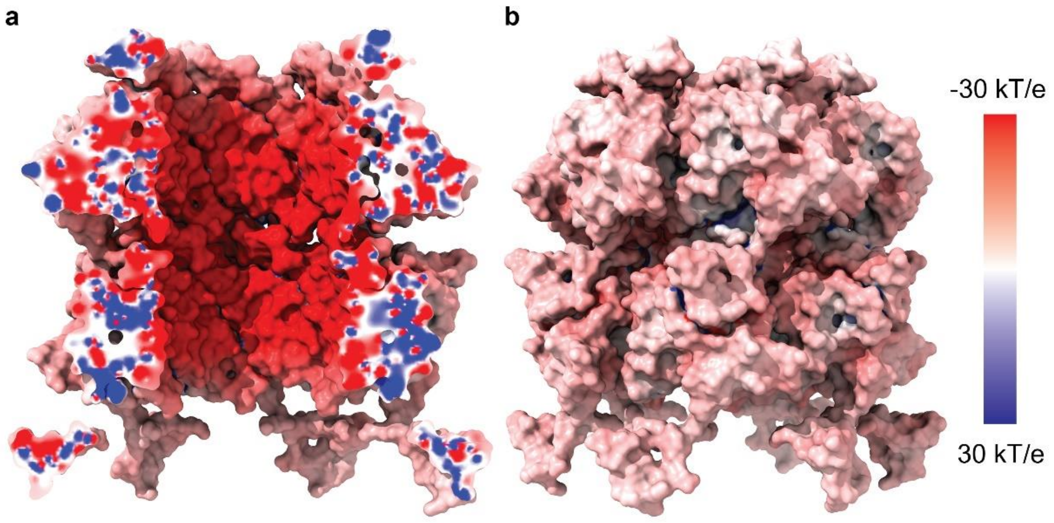

**Fig. S4.** Electrostatic potential of two rings of the SPP1 phage tail-tube. The structure (PDB ID 6YQ5) was prepared with PDB2PQR(*52*) and the potential was computed with the Adaptive Poisson-Boltzmann Solver (APBS) program(*53*). The tail features generally a negative electrostatic surface – especially the highly negatively charged lumen. The tail-tube is shown in a side view from the a) inside and b) outside.

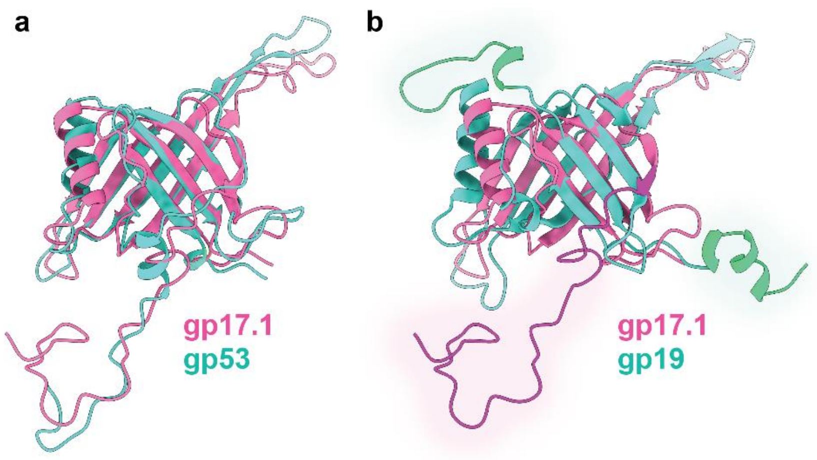

**Fig. S5.** Structural alignment of the TTPs gp17.1 (*Siphoviridae*, SPP1 phage) and gp53(*10*) (*Siphoviridae*, 80$\alpha$ phage), gp19(*54*) (*Myoviridae*, T4 phage). Structural alignments were performed with ChimeraX using the Needleman-Wunsch algorithm with a BLOSUM-62 residue similarity matrix of weight 0.7 and secondary structure scoring of weight 0.3.(*50*) In general, all TTPs share a common fold with RMSDs of 0.97 Å for gp17.1 and gp53 (**a**, between 67 pruned CA atom pairs), and 1.36 Å for gp17.1 and gp19 (**b**, between 21 pruned CA atom pairs. Structural differences are highlighted in magenta for gp17.1 and green for gp19. All TTPs share a common fold consisting of a $\beta$-sandwich-type fold that is flanked by an $\alpha$-helix. gp17.1 and gp53 additionally feature similar loop regions – the C-arm (143-176) and the loop (40-59). gp19 lacks a C-terminal extension but has two additional loop regions.

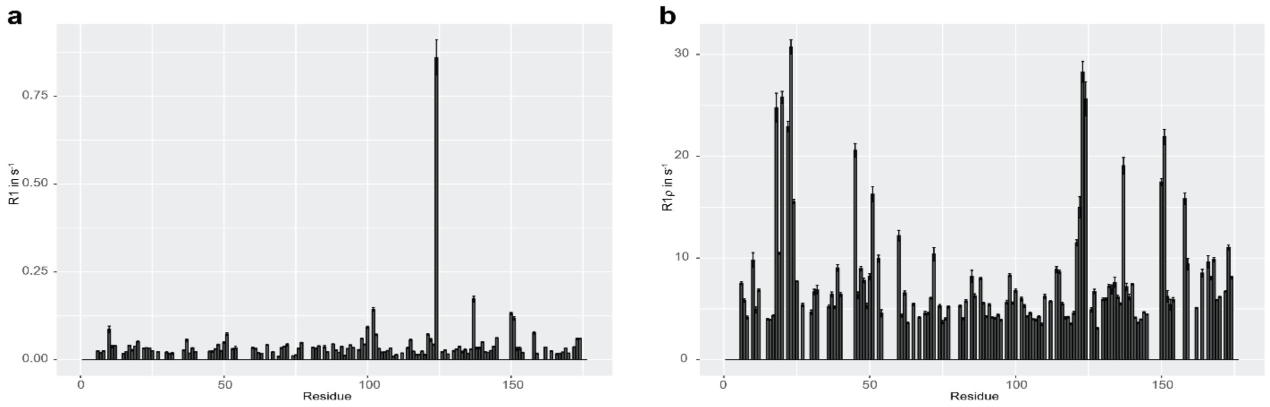

**Fig. S6.** Relaxation rates a) R1 and b) R1ρ as a function of residue number. Missing values represent residues that superimpose in the 2D hNH spectrum. The data were collected at 40 kHz MAS, 900 MHz external magnetic field strength and a temperature of +18 °C. The R1ρ rates were collected with a spin lock field of 5 kHz.

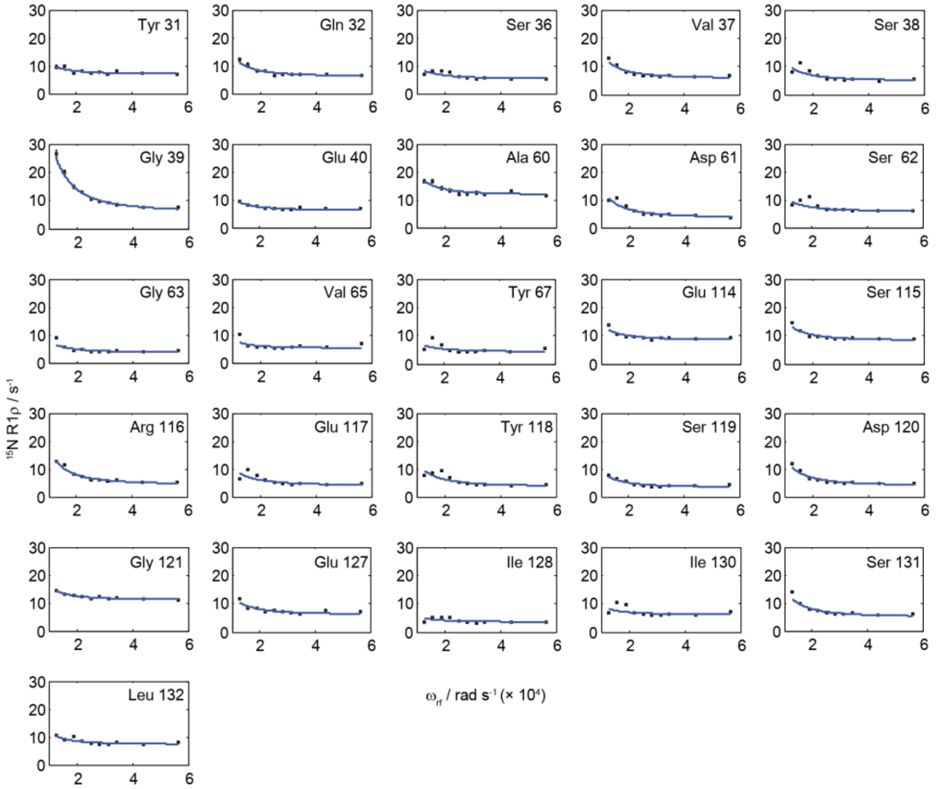

**Fig. S7.** Combined fit of $^{15}$N R1ρ relaxation dispersion plots of residues belonging to the inner β-barrel to a two-state Bloch McConell exchange process. The fit was conducted with a global exchange coefficient $k_{ex}$, individual $\phi_{ex}$ and individual R1ρ rates. The data were collected at 40 kHz MAS, 900 MHz external magnetic field strength and a temperature of +18 °C.

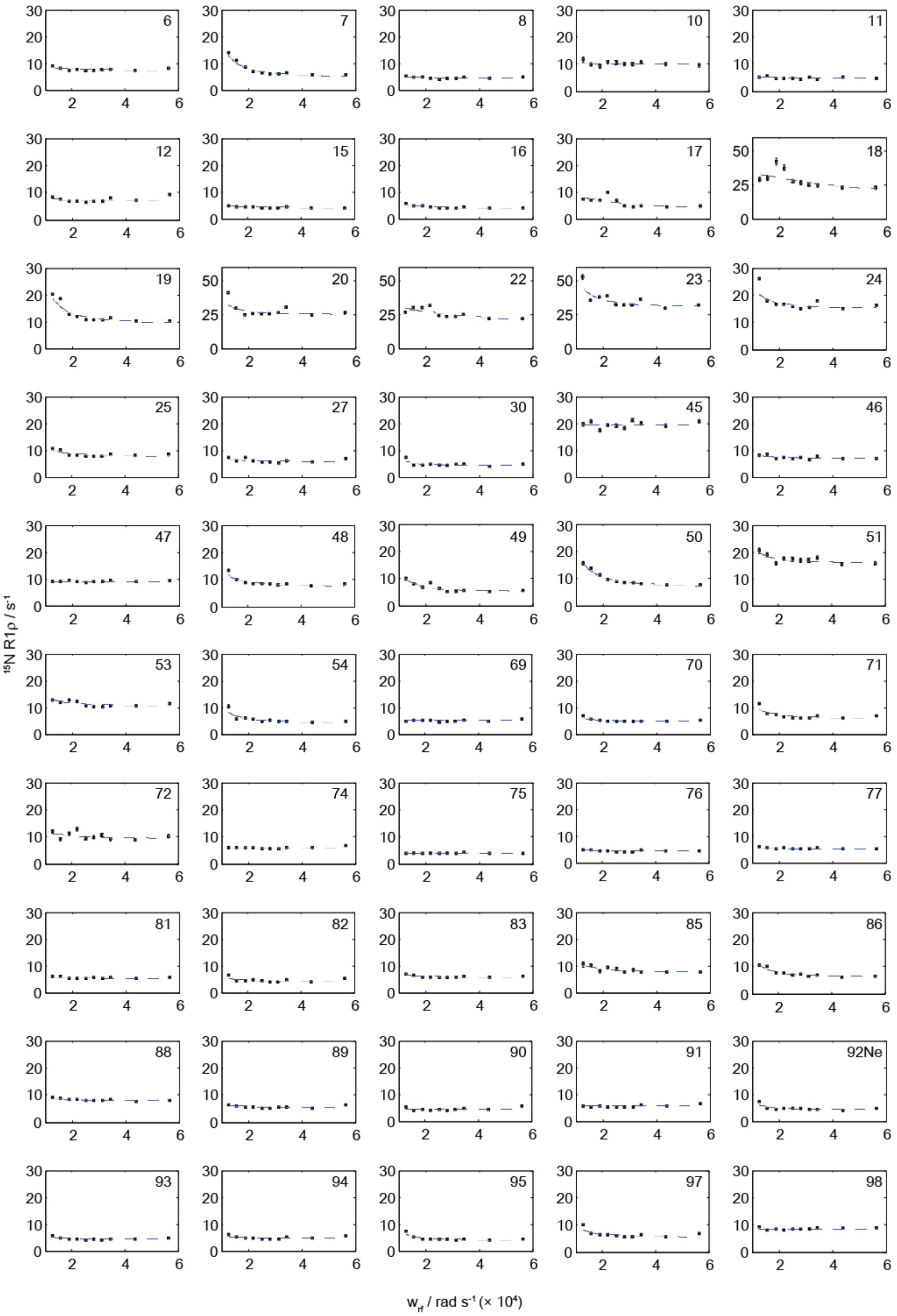

**Fig. S8.** $^{15}N$ R1ρ relaxation dispersion plots of residues 6 to 98 of gp17.1. The line represents the best fit to a two-state Bloch McConell equation as described in the Methods. Missing residues superimpose in the 2D hNH spectrum and/or are unassigned. The data were collected at 40 kHz MAS, 900 MHz external magnetic field strength and a temperature of +18 °C. Ile 18, Ala 20, Thr 22 and Gly 23 have a different y-axis scaling.

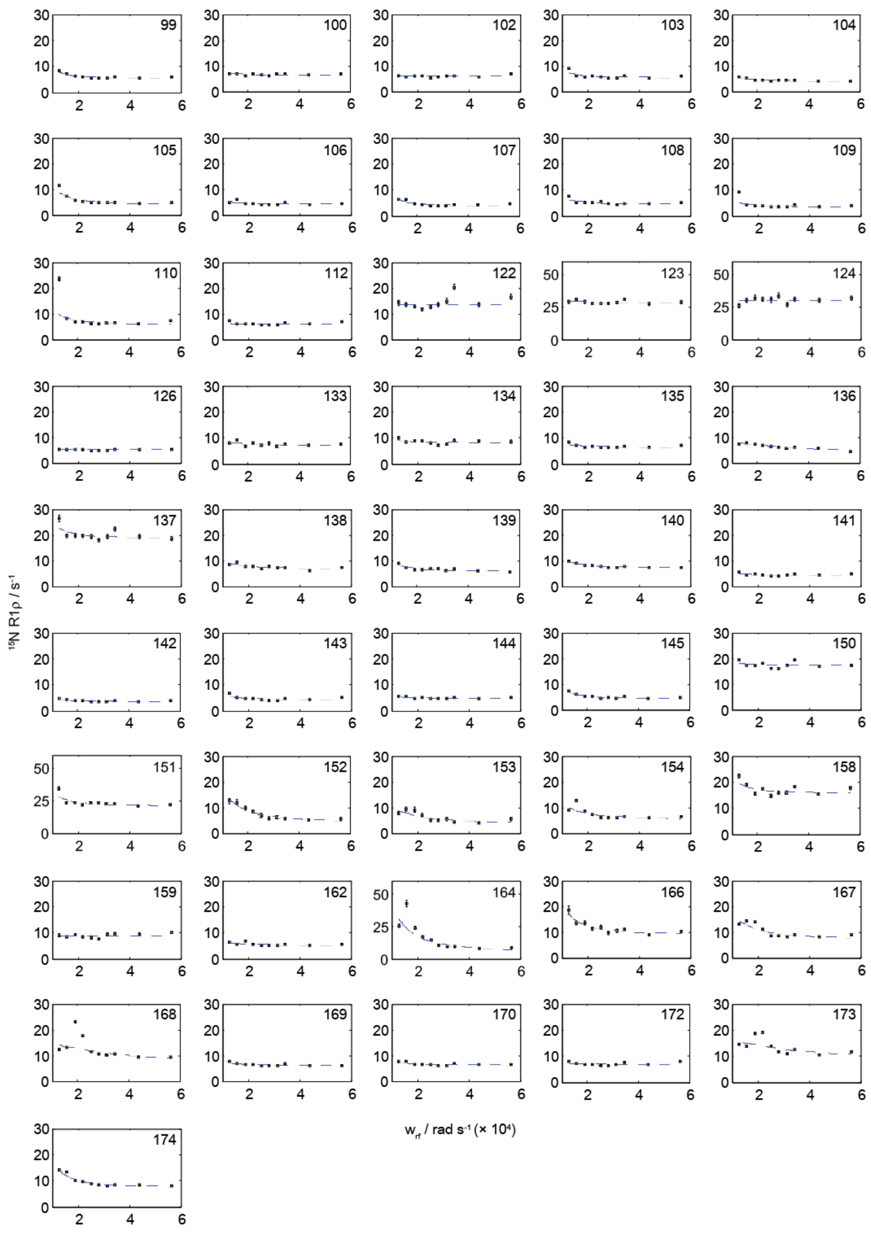

**Fig. S9.** $^{15}$N R1ρ relaxation dispersion plots of residues 99 to 174 of gp17.1. The line represents the best fit to a two-state Bloch McConell equation as described in the Methods. Missing residues superimpose in the 2D hNH spectrum and/or are unassigned. The data were collected at 40 kHz MAS, 900 MHz external magnetic field strength and a temperature of +18 °C. Glu 123, Gly 124, Val 151 and Gly 164 have a different y-axis scaling.

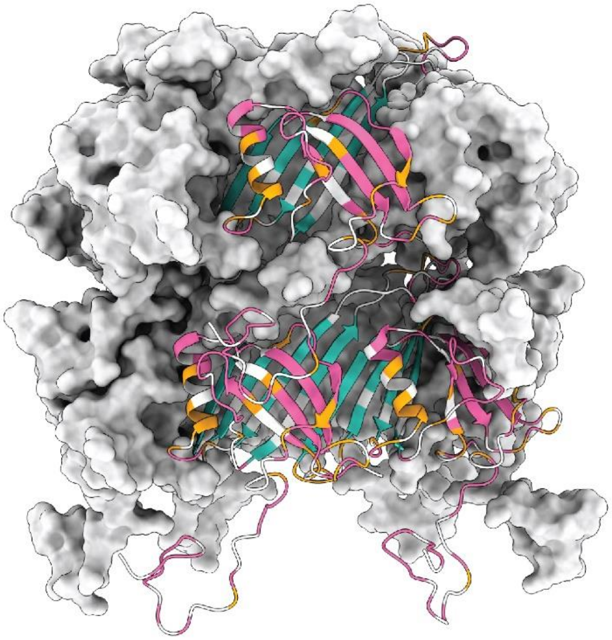

**Fig. S10.** $^{15}N$ relaxation dispersion mapped onto three subunits of two rings of the SPP1 tail tube. Residues highlighted in pink and turquoise show non-flat relaxation dispersion profiles and are, thus, involved in motions on the millisecond timescale. Residues turquoise can bet fitted in a combined manner to a two-state Bloch McConell exchange process. Residues in orange show flat profiles, residues in white superimpose or are unassigned.

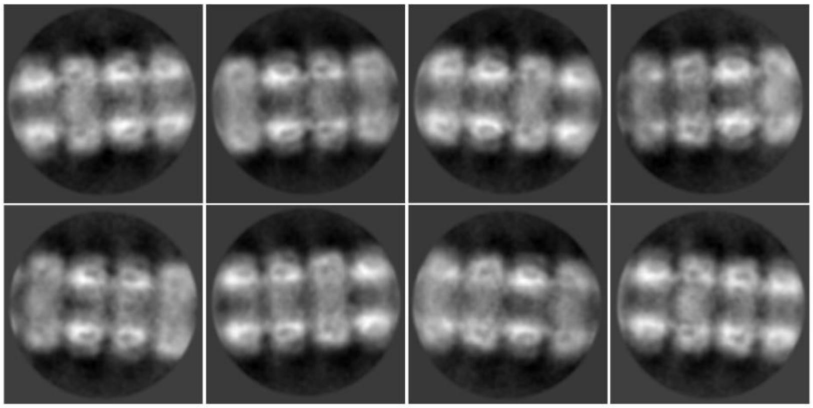

**Fig. S11.** Exemplary 2D class averages for the straight segments.

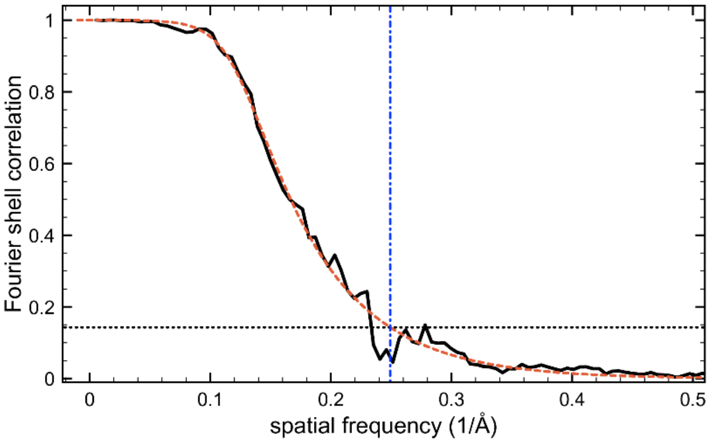

**Fig. S12.** Fourier shell correlation (FSC) calculated between two half maps. According to the 0.143 criterion the obtained resolution is 4.0 Å. The FSC curve was fitted using $1/[e^{((x-A)/B)}+1]^{C}$ , yielding A=0.122, B=0.015, and C=0.228 .

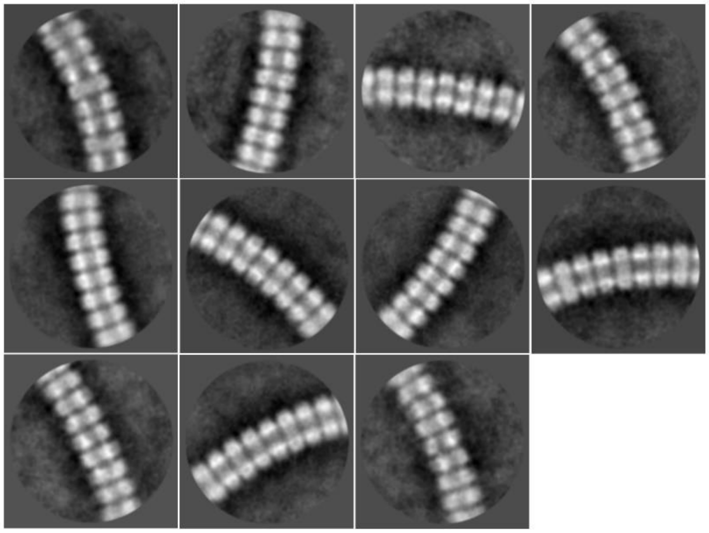

**Fig. S13.** 2D class averages of bent tail tubes.

**Table S1.** Methyl-labeled and/or deuterated protein samples used in this study. The precursors were supplemented to the bacterial culture as described by us in Zinke et al. 2018 and in the protocols provided by NMR-Bio (Grenoble).

| Labeling scheme | Labeling alias | Methyl-precursor |
|---|---|---|
| u-($^{2}$H, $^{13}$C, $^{15}$N) | uniform | n/a |
| isoleucine-($^{1}$Hδ1, $^{13}$Cδ1)-u-$^{15}$N | isoleucine-methyl | 2-Ketobutyric acid-4-$^{13}$C-3,3-$d_2$ (Sigma Aldrich)(*55*) |
| 50% isoleucine-($^{1}$Hδ1, $^{13}$Cδ1)-u-$^{14}$N & 50% u-$^{15}$N | isoleucine-methyl mix | 2-Ketobutyric acid-4-$^{13}$C-3,3-$d_2$ (Sigma Aldrich)(*55*) |
| alanine-($^{1}$Hβ, $^{13}$Cβ)-u-$^{15}$N | alanine-methyl | SLAM-$A^{\beta}$ Kit from NMR-Bio(*56*) |
| 50% alanine-($^{1}$Hβ, $^{13}$Cβ)-u-$^{14}$N & 50% u-$^{15}$N | alanine-methyl mix | SLAM-$A^{\beta}$ Kit from NMR-Bio(*56*) |
| leucine-($^{1}$Hδ, $^{13}$Cδ)$^{proR}$-valine-($^{1}$Hγ, $^{13}$Cγ)$^{proR}$-u-$^{15}$N | LV-methyl | DLAM-$LV^{proR}$ Kit from NMR-Bio(*57*) |
| 50% leucine-($^{1}$Hδ, $^{13}$Cδ)$^{proR}$- valine-($^{1}$Hγ, $^{13}$Cγ)$^{proR}$-u-$^{14}$N & 50% u-$^{15}$N | LV-methyl mix | DLAM-$LV^{proR}$ Kit from NMR-Bio(*57*) |
| threonine-($^{1}$Hγ2, $^{13}$Cγ2)-u-$^{15}$N | threonine-methyl | SLAM-T Kit from NMR-Bio(*58*) |
| 50% threonine-($^{1}$Hγ2, $^{13}$Cγ2)-u-$^{14}$N & 50% u-$^{15}$N | threonine-methyl mix | SLAM-T Kit from NMR-Bio(*58*) |
| methionine-($^{1}$Hε, $^{13}$Cε)-u-$^{15}$N | methionine methyl | SLAM-$M^{\varepsilon}$ Kit from NMR-Bio(*59*) |
| alanine-($^{1}$Hβ, $^{13}$Cβ)-isoleucine-($^{1}$Hδ1, $^{13}$Cδ1)-( $^{1}$Hγ2, $^{13}$Cγ2)-leucine-($^{1}$Hδ, $^{13}$Cδ)$^{proR/proS}$-valine-($^{1}$Hγ, $^{13}$Cγ)$^{proR/proS}$- u-$^{15}$N | assignment | QLAM-$A^{\beta}I^{\delta1/\gamma2}LV^{proR/proS}$ Kit from NMR-Bio(*60*) |

**Table S2.** Summary of the acquired solid-state NMR spectra including their purpose. Methyl-labeled and/or deuterated samples were studied at 40 kHz MAS. The fully-protonated samples were studied at 11 kHz MAS.

| | | | Summary of NMR spectra used in this study | |
|---|---|---|---|---|
| **Spinning Speed (rotor type)** | **Protein Sequence** | **Experiments** | **Sample** | **Purpose** |
| 11 kHz (3.2 mm) | gp17.1 | 2D hCC | fully-protonated | fingerprint |
| | ∆N-3 gp17.1 | 2D hCC | fully-protonated | fingerprint |
| 40 kHz (1.9 mm) | gp17.1 | 2D hCH | isoleucine-methyl | fingerprint |
| | | | alanine-methyl | |
| | | | LV-methyl | |
| | | | threonine-methyl | |
| | | | methionine methyl | |
| | | | assignment | |
| | | 3D HNhH | isoleucine-methyl | long-distance restraints |
| | | | alanine-methyl | |
| | | | LV-methyl | |
| | | | threonine-methyl | |
| | | | methionine methyl | |
| | | | alanine-methyl mix | intermolecular long-distance restraints |
| | | | isoleucine-methyl mix | |
| | | | LV-methyl mix | |
| | | | threonine-methyl mix | |
| | | 3D HChH | isoleucine-methyl | long-distance restraints |
| | | | LV-methyl | |
| | | 4D HNhhNH | uniform | long-distance restraints |
| | | pseudo-3D | uniform | relaxation rates |

**Table S3.** Parameters used for the spectral reconstruction of the non-uniformly sampled 4D spectrum. The reconstruction was conducted using the hmsIST software package.(*61*)

| | **Complex points after reconstruction** | | | | |
|---|---|---|---|---|---|
| **Experiment** | **F1** | **F2** | **F3** | **Iterations** | **Level Multiplier** |
| 4D HNhhNH | 48 | 30 | 48 | 3000 | 0.995 |

**Table S4.** Pulse program parameters. All experiments were conducted at a magic-angle spinning rate of 40 kHz and an external $B_0$ field corresponding to 900 MHz $^1$H Larmor frequency. Unless mentioned otherwise, carrier positions were set to the center of the chemical shift range.

| **Parameter** | **Value** | | | |
|---|---|---|---|---|
| **Experiment** | **2D hCH** | **3D HNhH** | **3D HChH** | **4D HNhhNH** |
| | | **Samples** | | |
| Labeling alias | isoleucine-methyl<br>alanine-methyl<br>LV-methyl<br>threonine-methyl<br>methionine methyl | isoleucine-methyl<br>isoleucine-methyl mix<br>alanine-methyl<br>alanine-methyl mix<br>LV-methyl<br>LV-methyl mix<br>threonine-methyl<br>threonine-methyl mix<br>methionine methyl | isoleucine-methyl<br>LV-methyl | uniform |
| | | **Recycle delay** | | |
| Recycle delay | 1 s | 1.15 s | 1 s | 1 s |
| | | **90° initial $^1$H excitation pulse** | | |
| R.f. power | 100 kHz | 100 kHz | 100 kHz | 83.3 kHz |
| Duration | 2.5 µs | 2.5 µs | 2.5 µs | 3 µs |
| Carrier position | | 8.5 ppm | | 8.5 ppm |
| | | **$^1$H evolution time** | | |
| WALTZ r.f. power | | 3.4 kHz ($^{15}$N) | 4.9 kHz ($^{13}$C) | 7.9 kHz ($^{15}$N) |
| WALTZ pulse duration | | 60 µs | 60 µs | 60 µs |
| WALTZ carrier position | | 117.7 ppm | | 117.7 ppm |
| | | **$^1$H-$^{15}$N CP step** | | |
| $^1$H r.f. power | | 81 kHz | | 78.7 kHz |
| $^1$H carrier position | | 8.5 ppm | | 8.5 ppm |
| $^{15}$N r.f. power | | 29.6 kHz | | 30.4 kHz |
| $^{15}$N carrier position | | 117.7 ppm | | 117.7 ppm |
| Ramp shape | | Ramp 80-100% on $^1$H | | Ramp 80-100% on $^1$H |
| Duration | | 1400 µs | | 900 µs |
| | | **$^{15}$N evolution time** | | |
| WALTZ r.f. power | | 9.5 kHz ($^1$H) | | 8.5 kHz ($^1$H) |
| WALTZ pulse duration | | 40 µs | | 50 µs |
| WALTZ carrier position | | 8.5 ppm | | 8.5 ppm |
| | | **$^1$H-$^{13}$C CP step** | | |
| $^1$H r.f. power | 52.5 kHz | | 54.0 kHz | |
| $^1$H carrier position | | | | |
| $^{13}$C r.f. power | 10.2 kHz | | 10.2 kHz | |
| $^{13}$C carrier position | | | | |
| Ramp shape | Ramp 80-100% on $^1$H | | Ramp 80-100% on $^1$H | |
| Duration | 2 ms | | 1.7 ms | |
| | | **$^{13}$C evolution time** | | |
| WALTZ r.f. power | 2.8 kHz ($^1$H) | | 3.6 kHz ($^1$H) | |
| WALTZ pulse duration | 40 µs | | 40 µs | |
| WALTZ carrier position | | | | |
| | | **90° $^{15}$N/$^{13}$C flip pulses** | | |
| R.f. power | 50 kHz | 35.7 kHz | 50 kHz | 35.7 kHz |
| Duration | 5 µs | 7 µs | 5 µs | 7 µs |
| Carrier position | | 117.7 ppm | | 117.7 ppm |
| | | **Water suppression** | | |
| T delay | 44 ms | 44 ms | 44 ms | 42 ms |
| Spoil pulse r.f. power | 43.6 kHz | 43.6 kHz | 43.6 kHz | |
| Spoil pulse duration | 1 ms | 1 ms | 1 ms | |
| Spoil pulse shape | Ramp 100-60% | Ramp 100-60% | Ramp 100-60% | |
| First pulse duration in train | 33 ms | 33 ms | 33 ms | 38 ms |
| Second pulse duration in train | 56 ms | 56 ms | 56 ms | 64.4 ms |
| Train r.f. power | 13.8 kHz | 13.8 kHz | 13.8 kHz | 16.4 kHz |

| | | | | |
|---|---|---|---|---|
| Wat. sup. carrier position | 4.9 ppm | 4.9 ppm | 4.9 ppm | 4.9 ppm |
| Loops through train (n) | 1 | 1 | 1 | 1 |
| Gradient pulse shape | | | | SINE.100 |
| Gradient pulse power | | | | 100% |
| Gradient pulse duration | | | | 1 ms |
| Delay for ring down | | | | 500 µs |
| **$^{15}$N-$^{1}$H CP step** | | | | |
| $^{1}$H r.f. power | | 81.6 kHz | | 78.7 kHz |
| $^{1}$H carrier position | | 8.5 ppm | | 8.5 ppm |
| $^{15}$N r.f. power | | 33.5 kHz | | 31.1 kHz |
| $^{15}$N carrier position | | 117.7 ppm | | 117.7 ppm |
| Ramp shape | | Ramp 100-80% on $^{1}$H | | Ramp 100-80% on $^{1}$H |
| Duration | | 900 µs | | 800 µs |
| **$^{13}$C-$^{1}$H CP step** | | | | |
| $^{1}$H r.f. power | 52.5 kHz | | 51.6 kHz | |
| $^{1}$H carrier position | | | | |
| $^{13}$C r.f. power | 10.2 kHz | | 10.2 kHz | |
| $^{13}$C carrier position | | | | |
| Ramp shape | Ramp 100-80% on $^{1}$H | | Ramp 100-80% on $^{1}$H | |
| Duration | 2 ms | | 900 µs | |
| **$^{1}$H-$^{1}$H RFDR Mixing** | | | | |
| RFDR r.f. power | | 100 kHz | 100 kHz | 100 kHz |
| RFDR mixing time | | 10 ms | 6 ms | 6 ms |
| **2nd $^{1}$H-$^{15}$N CP step** | | | | |
| $^{1}$H r.f. power | | | | 78.7 kHz |
| $^{1}$H carrier position | | | | 8.5 ppm |
| $^{15}$N r.f. power | | | | 31.1 kHz |
| $^{15}$N carrier position | | | | 117.7 ppm |
| Ramp shape | | | | Ramp 80-100% on $^{1}$H |
| Duration | | | | 800 µs |
| **Acquisition** | | | | |
| $^{15}$N WALTZ r.f. power | | 3.4 kHz | | 7.9 kHz |
| $^{15}$N WALTZ pulse duration | | 60 µs | | 60 µs |
| $^{15}$N WALTZ carrier position | | 117.7 ppm | | 117.7 ppm |
| $^{13}$C WALTZ r.f. power | 9.8 kHz | 5 kHz | 4.9 kHz | |
| $^{13}$C WALTZ pulse duration | 60 µs | 60 µs | 60 µs | |
| $^{13}$C WALTZ carrier position | | 24.9 ppm | | |

**Table S5.** Acquisition parameters for 2D, 3D, and 4D spectra. The highest dimension is always the direct dimension. Parameters for 2D hCH, 3D HNhH, and 3D HChH spectra are exemplarily shown for the LV-methyl labeled species. Acquired points for further methyl-labeled samples vary due to inherently different chemical shift dispersions. The 4D HNhhNH spectrum was recorded on the uniformly labeled sample and with non-uniform sampling (25 %).

| | Acquisition time / ms (number of complex points) | | | | | | |
|---|---|---|---|---|---|---|---|
| **Experiment** | **F1** | **F2** | **F3** | **F4** | **ns** | **Total Number of acquired points** | **Total Time** |
| 2D hCH (LV-methyl) | 15.2 ms (38) ($^{13}$C) | 20 ms (512) ($^{1}$H) | N/A | N/A | 48 | 76 | 1 h 23 min |
| 3D HNhH (LV-methyl) | 7.5 ms (30) ($^{1}$H) | 15 ms (45) ($^{15}$N) | 20 ms (512) ($^{1}$H) | N/A | 16 | 5400 | 1 d 15 h 34 min |
| 3D HChH (LV-methyl) | 7 ms (14) ($^{1}$H) | 10 ms (21) ($^{13}$C) | 20 ms (512) ($^{1}$H) | N/A | 32 | 1176 | 15 h 41 min |
| 4D HNhhNH | 10.6 ms (32) ($^{15}$N) | 6.2 ms (23) ($^{1}$H) | 10.6 ms (32) ($^{15}$N) | 21.3 ms (512) ($^{1}$H) | 8 | 47448/188416 | 7 d 2 h 28 min |

**Table S6.** Processing parameters for 2D, 3D and 4D spectra. The highest dimension is always the direct dimension.

| | Points after FT | | | | Window function | | | |
|---|---|---|---|---|---|---|---|---|
| **Experiment** | **F1** | **F2** | **F3** | **F4** | **F1** | **F2** | **F3** | **F4** |
| 2D hCH | 1k ($^{13}C$) | 4k ($^{1}H$) | N/A | N/A | $sin^2$, φ=45° | $sin^2$, φ=45° | N/A | N/A |
| 3D HNhH | 128 ($^{1}H$) | 128 ($^{15}N$) | 4k ($^{1}H$) | N/A | $sin^2$, φ=60° | $sin^2$, φ=60° | $sin^2$, φ=60° | N/A |
| 3D HChH | 128 ($^{1}H$) | 128 ($^{13}C$) | 2k ($^{1}H$) | N/A | $sin^2$, φ=60° | $sin^2$, φ=60° | $sin^2$, φ=60° | N/A |
| 4D HNhhNH | 64 ($^{15}N$) | 64 ($^{1}H$) | 64 ($^{15}N$) | 1k ($^{1}H$) | $sin^2$, φ=90° | $sin^2$, φ=90° | $sin^2$, φ=90° | $sin^2$, φ=60° |

**Table S7.** Values for the fitting of the relaxation dispersion profiles of the inner β-barrel residues to a two-state Bloch McConell exchange process. Exemplary chemical shift differences between the two sites $p_A$ (95%) and $p_B$ (5%) are extracted from the $\phi_{ex}$ individual values.

| Residue | Global exchange coefficient $k_{ex}$ ($s^{-1}$) | Individual $\phi_{ex}$ ($rad^2\ s^{-2}$) | Individual $R_{1\rho}$ ($s^{-1}$) | Chemical shift differences (ppm) |
|---|---|---|---|---|
| 31 | 4528 ± 2349 | 105913 ± 89686 | 6.97 ± 0.39 | 2.64±2.23 |
| 32 | | 202516 ± 200301 | 6.10 ± 0.66 | 3.65±3.61 |
| 36 | | 120082 ± 93918 | 5.28 ± 0.20 | 2.81±2.19 |
| 37 | | 235173 ± 193440 | 5.75 ± 0.40 | 3.93±3.23 |
| 38 | | 184775 ± 140977 | 4.95 ± 0.21 | 3.48±2.66 |
| 39 | | 760021 ± 626743 | 5.90 ± 0.59 | 7.07±5.83 |
| 40 | | 118126 ± 84213 | 6.18 ± 0.27 | 2.78±1.98 |
| 60 | | 202920 ± 172788 | 11.67 ± 0.78 | 3.65±3.11 |
| 61 | | 288634 ± 234700 | 3.51 ± 0.27 | 4.35±3.54 |
| 62 | | 141308 ± 94503 | 5.79 ± 0.28 | 3.05±2.03 |
| 63 | | 111162 ± 89211 | 3.69 ± 0.13 | 2.7±2.17 |
| 65 | | 81359 ± 60740 | 5.21 ± 0.14 | 2.31±1.72 |
| 67 | | 93217 ± 65064 | 4.29 ± 0.12 | 2.47±1.72 |
| 114 | | 147325 ± 121743 | 8.43 ± 0.52 | 3.11±2.57 |
| 115 | | 196099 ± 15092 | 8.26 ± 0.24 | 3.59±2.91 |
| 116 | | 336711 ± 255809 | 4.37 ± 0.22 | 4.7±3.57 |
| 117 | | 180028 ± 133857 | 4.11 ± 0.19 | 3.44±2.55 |
| 118 | | 235362 ± 155622 | 3.74 ± 0.22 | 3.93±2.6 |
| 119 | | 153699 ± 131700 | 3.41 ± 0.19 | 3.18±2.72 |
| 120 | | 268443 ± 207292 | 4.02 ± 0.24 | 4.2±3.24 |
| 121 | | 131404 ± 108054 | 11,11 ± 0.39 | 2.94±2.41 |
| 127 | | 176669 ± 117660 | 6.19 ± 0.48 | 3.41±2.27 |
| 128 | | 52916 ± 43747 | 3.55 ± 0.14 | 1.86±1.54 |
| 130 | | 82325 ± 60274 | 6.13 ± 0.24 | 2.32±1.7 |
| 131 | | 265027 ± 202907 | 5.02 ± 0.22 | 4.17±3.19 |
| 132 | | 118696 ± 101386 | 7.30 ± 0.23 | 2.79±2.38 |

**Table S8.** Cryo-EM data collection, refinement and validation statistics

| | EMD-10792,<br>PDB ID 6YEG |
|---|---|
| **Data collection and processing** | |
| Magnification | 110,000 |
| Voltage (kV) | 200 |
| Total dose ($e^{-}/Å^2$) | 70 |
| Exposure time (s) | 3 |
| Movie frames (no.) | 120 |
| Defocus range (μm) | -0.2 to -1.7 |
| Pixel size (Å) | 0.935 |
| No. Micrographs | 855 |
| Symmetry imposed | C6, helical |
| Helical rise (Å) | 38.46 |
| Helical twist (º) | 21.89 |
| Final fibril images (no.) | 1866 |
| Final particle images (no.) | 5965 |
| Map resolution (Å) | 4.0 |
| FSC threshold | 0.143 |
| **Refinement** | |
| Initial density model used | *sphere model based on estimated helical symmetry* |
| Model composition | |
| Non-hydrogen atoms | 15768 |
| Protein residues | 2064 |
| Chains | 12 |
| R.m.s. deviations | |
| Bond lengths (Å) | 0.0063 |
| Bond angles (°) | 1.32 |
| **Validation** | |
| MolProbity score | 2.24 |
| Clashscore | 11.2 |
| Poor rotamers (%) | 0.8 |
| Ramachandran plot | |
| Favored (%) | 84.4 |
| Disallowed (%) | 0.3 |

**Table S9.** Overview long-range distance restraints from solid-state NMR.

| Categories | Sequence separation |
|---|---|
| intra-residue | 0 |
| sequential | 1 |
| medium-range | 2-4 |
| long-range | >4 |

| NMR inter (only between subunits) | Number |
|---|---|
| intra-residue | 0 |
| sequential | 0 |
| medium-range | 0 |
| long-range | 17 |

| NMR ambiguous (within and between subunits) | Number |
|---|---|
| intra-residue | 12 |
| sequential | 120 |
| medium-range | 75 |
| long-range | 480 |

| Hydrogen bonds (within subunit) | Number |
|---|---|
| intra-residue | 0 |
| sequential | 0 |
| medium-range | 16 |
| long-range | 80 |

| Hydrogen bonds (between subunits) | Number |
|---|---|
| intra-residue | 0 |
| sequential | 0 |
| medium-range | 0 |
| long-range | 12 |

**Table S10.** Violations of solid-state NMR long-range distance restraints.

| NMR inter (only between subunits) | Number |
|---|---|
| total number of restraints | 17 |
| violations 7.0 – 8.0 Å | 1 |
| violations 8.0 – 9.0 Å | 1 |
| violations 9.0 – 10.0 Å | 0 |
| violations > 10 Å | 0 |

| NMR ambiguous (within and between subunits) | Number |
|---|---|
| total number of restraints | 687 |
| violations 7.0 – 8.0 Å | 63 |
| violations 8.0 – 9.0 Å | 35 |
| violations 9.0 – 10.0 Å | 18 |
| violations > 10 Å | 9 |

**Table S11.** Ensemble RMSDs after hybrid structure calculation (PDB ID 6YQ5). For each pair from the ensemble the RMSD between coordinates of an atom selection was computed using the Kabsch algorithm. These values were then averaged over all combinations and the standard deviation of these was computed. There is significant difference between the RMSDs within a single subunit and the corresponding RMSD values computed for the entire assembly.

| **Subunit** | **RMSD** |
|---|---|
| C-alpha atoms | 1.1 ± 0.52 Å |
| Backbone atoms | 1.1 ± 0.5 Å |
| Heavy atoms | 1.8 ± 0.87 Å |

| **Full assembly (12-mer)** | **RMSD** |
|---|---|
| C-alpha atoms | 1.1 ± 0.52 Å |
| Backbone atoms | 1.1 ± 0.51 Å |
| Heavy atoms | 1.8 ± 0.87 Å |

**Movie S1.**

Structure of the tail tube of the bacteriophage SPP1. Structural features of the complex as well as the structural restraints from solid-state NMR and cryo-EM are visualized.

**Movie S2.**

Bending of the tail tube of the bacteriophage SPP1 is facilitated by stretching of certain linker regions.
